# Microglia-specific transduction via AAV11 armed with IBA1 promoter and miRNA-9 targeting sequences

**DOI:** 10.1101/2024.07.09.602653

**Authors:** Nengsong Luo, Kunzhang Lin, Yuxiang Cai, Xiaokai Sui, Zilian Zhang, Jiayong Xing, Gangning Liu, Wenjia Yuan, Jie Wang, Fuqiang Xu

## Abstract

Microglia, as resident immune cells in the central nervous system (CNS), are closely related to human health and the pathogenesis of various CNS diseases, making them compelling targets for therapeutic interventions. However, functional and therapeutic studies of microglia remain significant challenges largely due to the lack of tools capable of efficiently and specifically transducing microglia. Herein, we evaluated the specificity and efficiency of various adeno-associated virus (AAV) vectors armed with the mIBA1 promoter and miRNA-9 targeting sequences in transducing microglia within the caudate putamen (CPu) brain region, and found that AAV11 mediates more specific and efficient transduction of microglia. Subsequently, we further demonstrated that AAV11 also exhibits high transduction specificity for microglia across various brain areas and within the spinal cord. Finally, by reducing the injection dosage, we employed AAV11 for sparse labeling of microglia. This work provides a promising tool for advancing both the functional investigation and therapeutic targeting of microglia.

## Introduction

Microglia, as the primary immune cells of the CNS, play an indispensable role in maintaining brain health through their involvement in immune surveillance, synaptic pruning, neuroprotection, and response to injury or disease^1–4^. Microglial dysfunctions are a key contributor to CNS aging and many CNS diseases including Alzheimer’s disease, chronic pain, brain cancers, depression, memory loss, and more^5–11^. Moreover, genome-wide association studies have pinpointed genetic variants in immune-related genes as pivotal risk factors in the initiation and progression of diseases, some of which are highly expressed in microglia^12–15^. Therefore, gaining insights into microglial structures and functions is fundamental to advancing our understanding of brain function and disease, and it has the potential to lead to new therapeutic strategies for a wide range of neurological conditions.

Structural and functional studies of resident microglia necessitate effective measures for specific targeting and genetic manipulation. Currently, generating germline transgenic mouse models remains the primary approach for introducing specific genes into microglia^16^, a process that is notably laborious, time-consuming, and inefficient. Recombinant viral vectors have been developed as valuable tools to address these limitations, allowing sufficient payload expression in microglia^17–19^. Åkerblom and colleagues utilized a microRNA-9-regulated lentiviral vector to selectively target resident microglia in the adult rodent brain^20^, thereby achieving precise transgene expression within these cells, but lentiviruses have a risk of tumorigenesis due to random integration into the host genome^21^. Targeting microglia with recombinant adeno-associated viruses (rAAVs) is a strategic priority due to their distinct advantages, including minimal pathogenicity and long-term gene expression^22–24^. rAAV had improved microglial transduction *in vitro* by carrying microglia-specific promoters such as F4/80 and CD68, while the *in vivo* transduction efficiency remained very low^25, 26^. Lin et al. successfully developed AAV9 variants (termed AAV-MGs) through directed evolution, which mediate efficient *in vitro* and *in vivo* microglial transduction^19^. Nevertheless, AAV-MGs still transduced non-microglial cells, thus requiring the use in combination with microglia-specifc Cre-transgenic strains to ensure specific targeting and manipulation of microglia. Therefore, it is crucial to persist in screening versatile AAVs capable of specifically labeling microglia with high efficiency.

In this study, we evaluated the specificity and efficiency of diverse AAV vectors in transducing microglia within the CPu brain region, when regulated by mIBA1 promoter and miRNA-9 targeting sequences, and found that AAV11 mediates more specific and efficient transduction of microglia. Subsequently, we further demonstrated that AAV11 also exhibits high transduction specificity for microglia across various brain areas and within the spinal cord. Finally, by reducing the injection dosage, we employed AAV11 for sparse labeling of microglia. This work provides a promising tool for advancing both the functional investigation and therapeutic targeting of microglia.

## Results

### Screening process of AAV vectors for microglia-specific targeting

Microglia play a pivotal role in brain development, maintenance of brain health, and regulation of neural activity^27^. However, there are currently limited AAVs available for studying and understanding microglia, possibly due to immunogenicity or the insufficient affinity of existing AAVs for microglia. Therefore, exploring new AAVs involves screening natural AAVs or modifying existing ones to enhance transduction efficiency in microglia. In this study, we compared several natural serotypes of AAV carrying customized gene expression cassettes, including AAV9, AAVhu.32, AAV11, and AAVrh32.33, for their specificity and efficiency in microglial cell transduction. These cassettes include a microglia-specific promoter, green fluorescent protein gene, and four repeated complementary microRNA-9-targeting (miR-9.T) sequences, illustrated in Figure 1A. Subsequently, we produced viral vectors using triple plasmid transient transfection method. The packaged viruses were then injected into the brains of mice, and after three weeks, brain samples were perfused, imaged, and subjected to signal quantification and comparison, as shown in Figure 1B.

**Figure 1.**
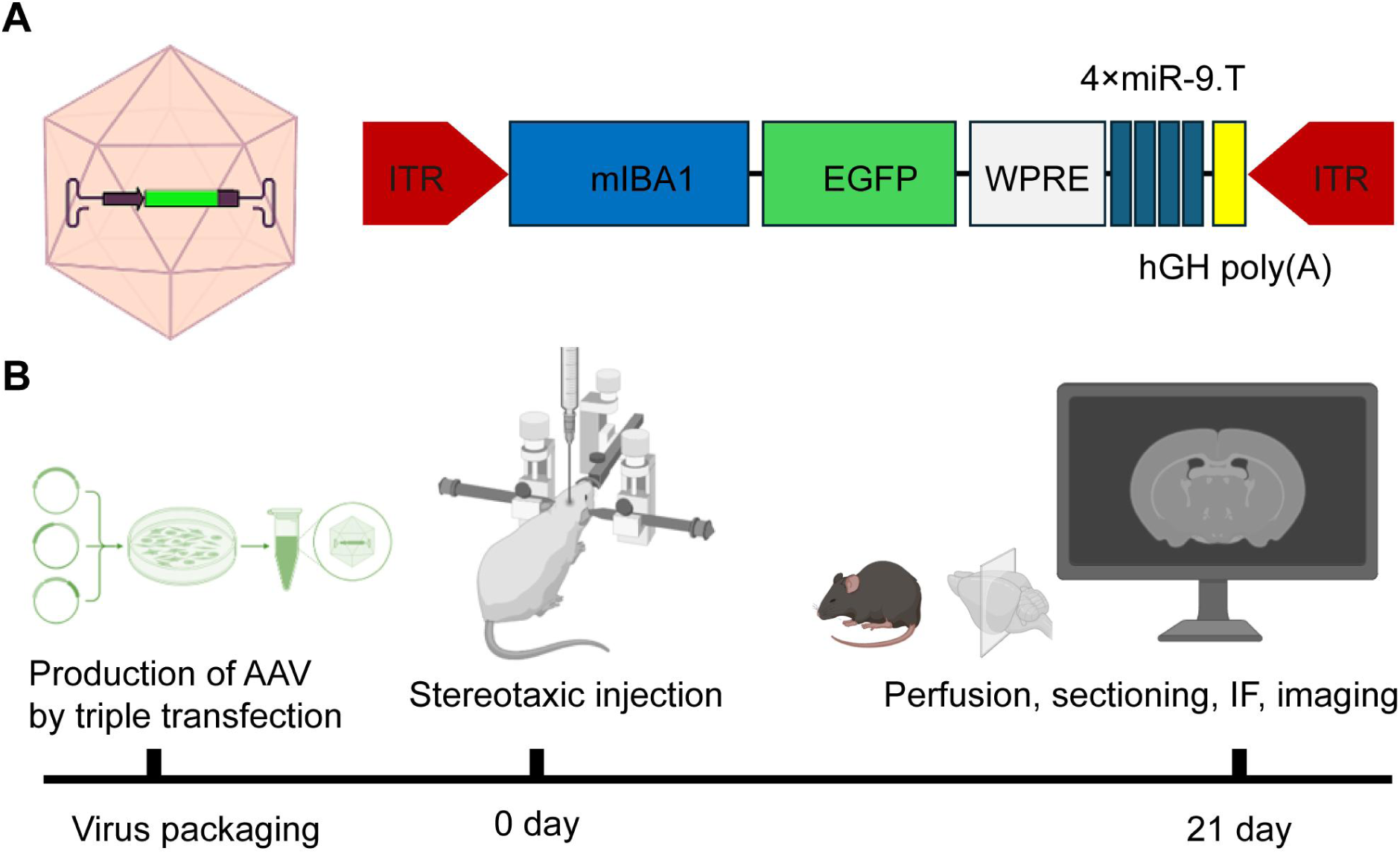
Flowchart illustrating the screening process for microglial-specific AAV vectors. A: AAV vectors comprise the constitutive IBA1 promoter followed by EGFP, WPRE, and polyA. Four repeated microRNA-9 (miR-9) targeting sequences (4×miR-9.T) were inserted between the WPRE and polyA sequences. B: Experimental process, including virus production, brain stereotactic injection, sampling, slicing, immunofluorescence staining and imaging.

### AAV11 specifically transduces striatum microglia with high efficiency

We focus on AAV mediated gene delivery systems for targeting microglia. We evaluated the specificity and efficiency of four different AAVs armed with the mIBA1 promoter and miRNA-9 targeting sequences in transducing microglia within the adult mouse brain. We separately injected the assembled AAVs into the CPu region at a dosage of 2 × 10^9^ vector genomes (VG) with a volume of 200 nL (Figure 2A). After three weeks, brain tissues were harvested and sliced, followed by immunofluorescence staining (Anti-IBA1), as shown in Figure 2B. To evaluate the cell-type specificity of these AAVs, we measured the colocalization of EGFP and microglial marker IBA1 (Figure 2C). To evaluate the transduction efficiency of AAVs on microglia, we compared the number of labeled microglia (Figure 2D). All these AAVs mediated robust EGFP expression in IBA1-positive cells, yet with varying levels of specificity and efficiency (Figure 2B-D). Quantification analysis of EGFP- and IBA1-double positive cells demonstrated that AAV11 infusion resulted in the highest degrees of specificity (approximately 98.9%, *p* < 0.0001, Figure 2C) and efficiency (approximately 947 cells, *p* < 0.0001, Figure 2D) for microglia within the striatum region.

**Figure 2.**
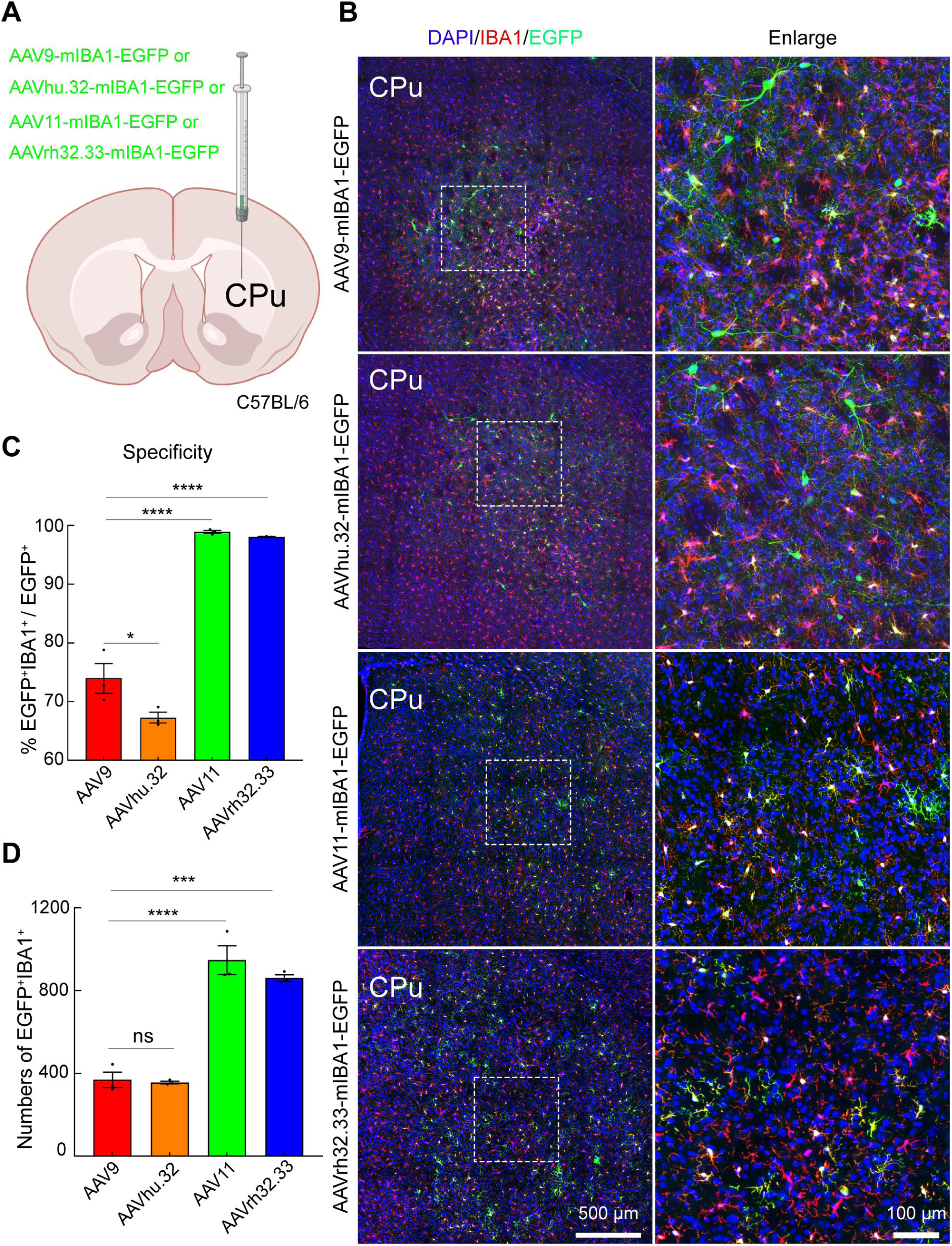
*In vivo* screens for microglia-targeting AAVs. A: Schematic diagram of virus injection. AAV9-mIBA1-EGFP, AAVhu.32-mIBA1-EGFP, AAV11-mIBA1-EGFP and AAVrh32.33-mIBA1-EGFP viruses (2 × 10^9^ VG each virus) were individually injected into striatum area of C57BL/6 mice. B: Representative confocal images showing marker expression for the indicated AAVs. Scale bar = 500 μm (left 4 panels), 100 μm (right 4 panels). C: Quantification of microglial transduction specificity in the striatum regions injected with the indicated AAVs (mean ± SEM; *n* = 3 mice per group). D: Quantification of microglial transduction efficiency in the striatum regions injected with the indicated AAVs (mean ± SEM; *n* = 3 mice per group). Statistical analyses were completed using one-way ANOVA followed by Tukey’s multiple comparisons test with alpha value of 0.05. ns, no significant difference; *, *p* < 0.05; ***, *p* < 0.001; ****, *p* < 0.0001.

### AAV11 specifically transduces microglia in DG region of mouse brain

The hippocampus is closely associated with learning and memory, as well as neurogenesis, emotional regulation, synaptic plasticity, and various diseases^28–30^. Recent studies have shown that various activities in the dentate gyrus (DG) of the hippocampus are associated with microglia^31–34^. We aim to investigate whether AAV11 can efficiently transduce microglia in the DG brain region. AAV11 were injected into the DG regions of the brain, with a dose of 2 × 10^9^ VG (Figure 3A). After three weeks, brain samples were perfused and subjected to immunofluorescence staining (Anti-IBA1). AAV11 demonstrated robust transduction capabilities in microglia within the hippocampal DG region (Figure 3B and 3C). Quantification analysis of EGFP- and IBA1-double positive cells demonstrated that AAV11 predominantly targeted microglia with approximately 97% specificity (Figure 3D).

**Figure 3.**
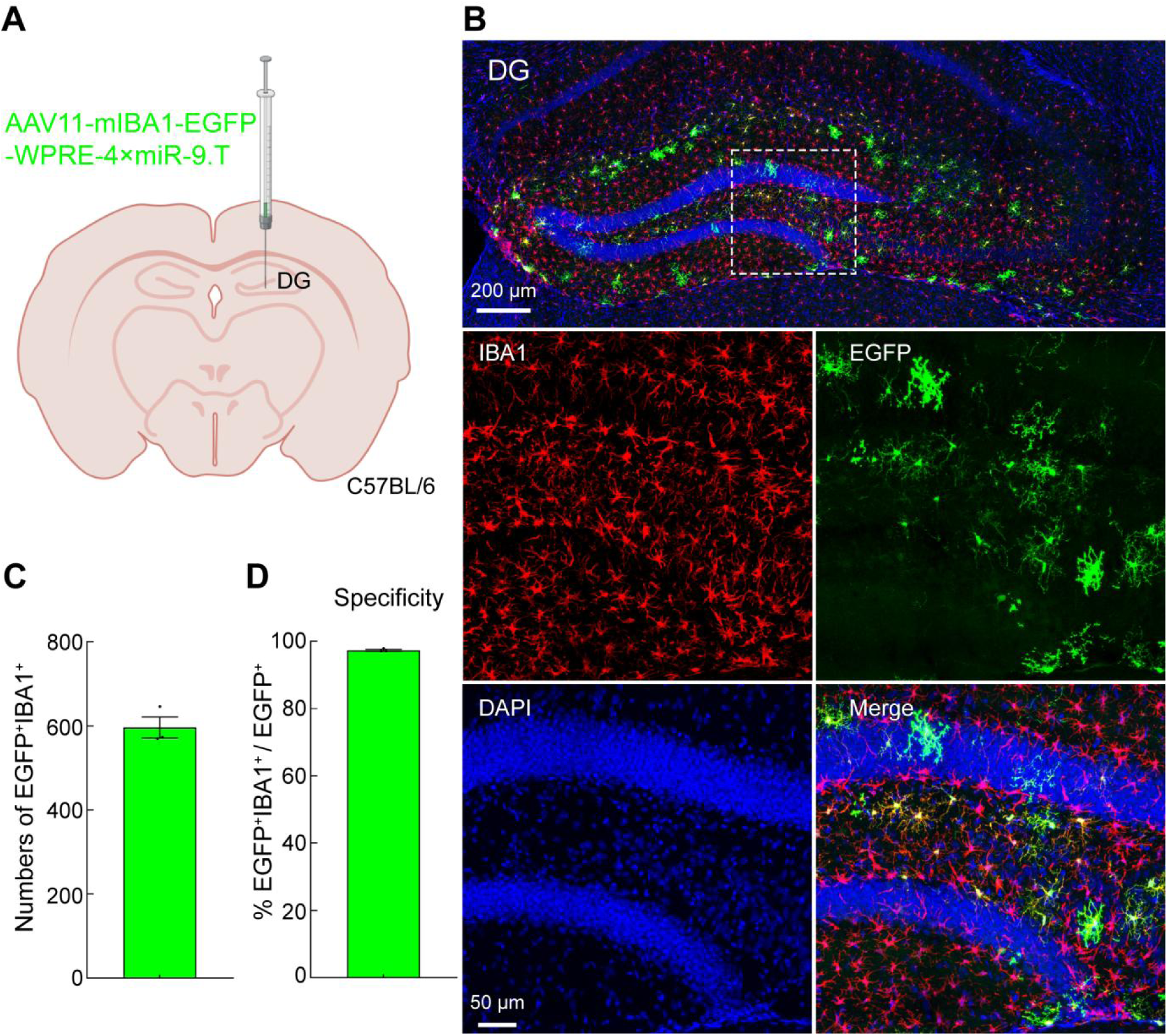
AAV11 efficiently and specifically transduces microglia in the DG brain region. A: Schematic diagram of virus injection. AAV11-mIBA1-EGFP viruses (2 × 10^9^ VG) were injected into DG area of C57BL/6 mice. B: Representative confocal images showing marker expression for the indicated AAVs. Scale bar = 200 μm (top 1 panel), 50 μm (bottom 4 panels). C: Quantifications showing microglial transduction efficiency for AAV11 (mean ± SEM; *n* = 3 mice per group). D: Quantifications showing microglial transduction specificity for AAV11 (mean ± SEM; *n* = 3 mice per group).

### AAV11 specifically transduces microglia in SNr region of mouse brain

We also injected AAV11 into the substantia nigra pars reticulata (SNr), a brain region primarily associated with motor regulation, posture control, and movement disorders, playing a crucial role in the study of basic neurophysiology and movement disorders^35, 36^. Previous studies have suggest that suppressing microglial activation in the SNr of PD model animals resulted in aggravated motor dysfunctions^37^. AAV11 were injected into the SNr with a dose of 2 × 10^9^ VG (Figure 4A). After three weeks, brain samples were perfused and subjected to immunofluorescence staining (Anti-IBA1). In the SNr, most microglia were observed to be labeled with green fluorescence (Figure 4B and 4C). Fluorescence colocalization analysis indicated that AAV11 mediated gene expression with approximately 90% specificity in the SNr microglia (Figure 4D).

**Figure 4.**
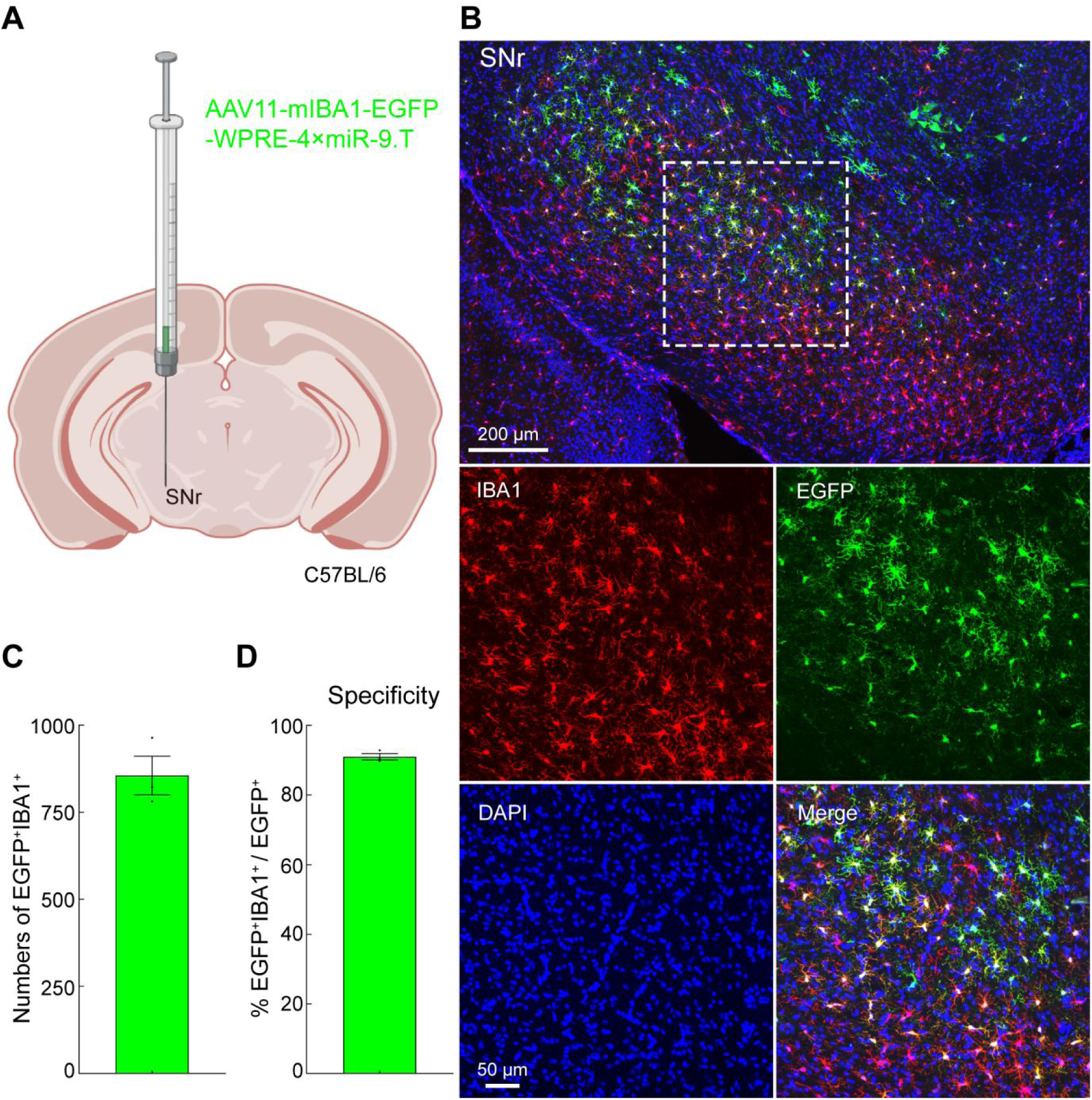
AAV11 efficiently and specifically transduces microglia in the SNr brain region. A: Schematic diagram of virus injection. AAV11-mIBA1-EGFP viruses (2 × 10^9^ VG) were injected into SNr area of C57BL/6 mice. B: Representative confocal images showing marker expression for the indicated AAVs. Scale bar = 200 μm (top 1 panel), 50 μm (bottom 4 panels). C: Quantifications showing microglial transduction efficiency for AAV11 (mean ± SEM; *n* = 3 mice per group). D: Quantifications showing microglial transduction specificity for AAV11 (mean ± SEM; *n* = 3 mice per group).

### AAV11 specifically transduces microglia in SSp region of mouse brain

Previous studies have shown that AAV viral vectors have lower specificity in transduction of cortical microglia cells^38^. To evaluate the specificity of AAV11 in cortical microglial cell transduction, we injected AAV11 into the primary somatosensory area (SSp) (Figure 5A), a brain region rich in projection neurons that is closely associated with sensory processing of pain, motor regulation, and neural plasticity^39, 40^. Studies have shown that the microglia in the SSp brain region are closely related to its development^41^. Immunofluorescence staining revealed that a small portion of the fluorescent signals did not colocalize with IBA1-positive signals (Figure 5B and 5C). Fluorescent colocalization analysis indicated approximately 70% specificity of AAV11-mediated gene expression in microglia within the SSp region (Figure 5D).

**Figure 5.**
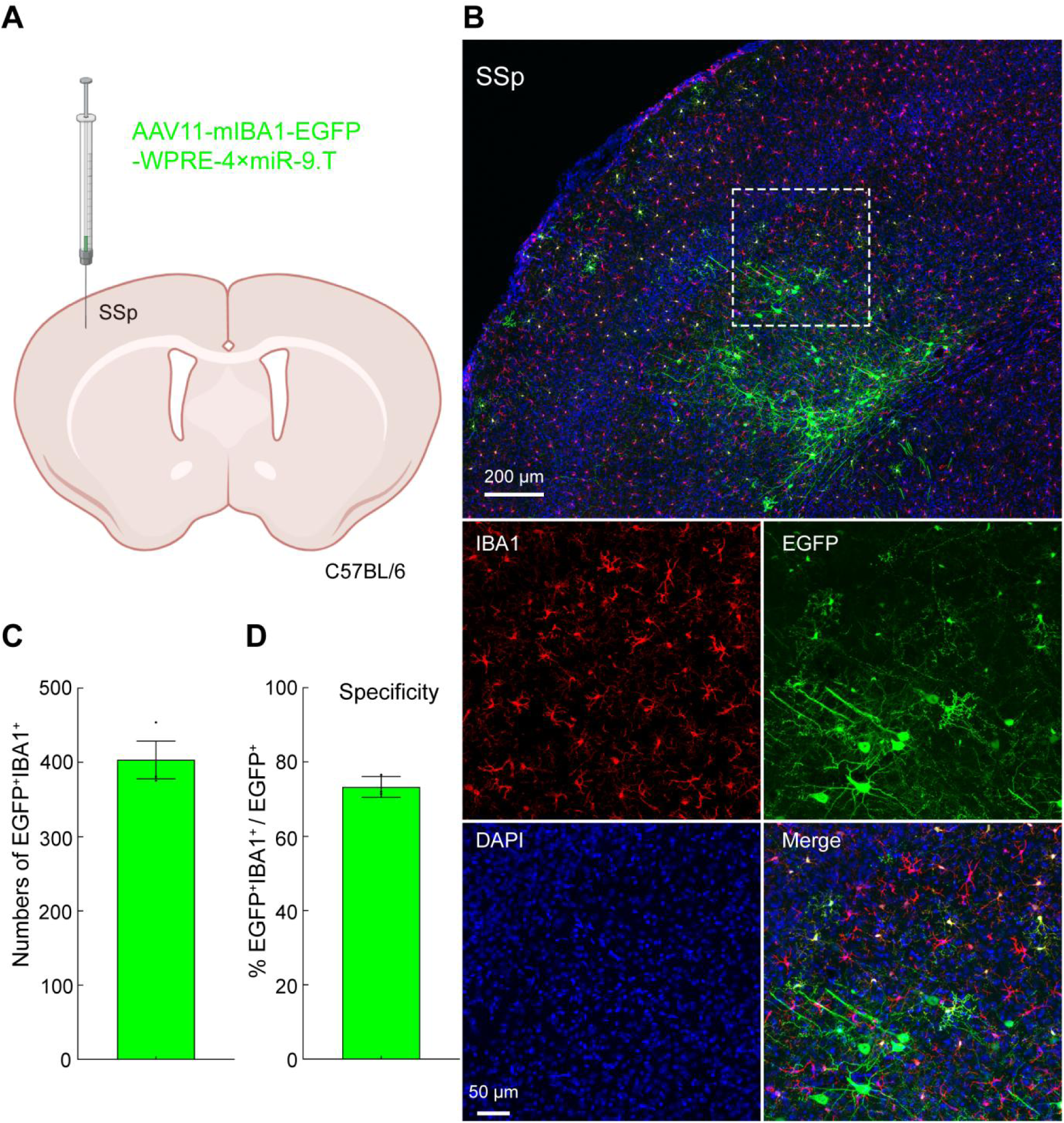
AAV11 efficiently and specifically transduces microglia in the SSp brain region. A: Schematic diagram of virus injection. AAV11-mIBA1-EGFP viruses (2 × 10^9^ VG) were injected into SSp area of C57BL/6 mice. B: Representative confocal images showing marker expression for the indicated AAVs. Scale bar = 200 μm (top 1 panel), 50 μm (bottom 4 panels). C: Quantifications showing microglial transduction efficiency for AAV11 (mean ± SEM; *n* = 3 mice per group). D: Quantifications showing microglial transduction specificity for AAV11 (mean ± SEM; *n* = 3 mice per group).

### AAV11 can specifically transduce microglia in the spinal cord

The spinal cord is the main pathway between the brain and the peripheral nervous system. In a healthy spinal cord, microglia are responsible for immune surveillance. However, when spinal cord injury occurs, the microenvironment changes dramatically, and microglia respond by producing both beneficial and detrimental effects on the injury^42^. Spinal cord microglia play an extremely important role in the repair of spinal cord injury, regulation of inflammatory response, generation of pain, and development of potential therapeutic strategies^43^. Therefore, it is necessary to develop gene delivery systems targeting spinal microglia. Here, we aim to investigate whether AAV11 can efficiently transduce microglia in the spinal cord. AAV11 were injected into the lumbar segment of the spinal cord, with a dose of 5 × 10^9^ VG (Figure 6A). After three weeks, brain samples were perfused and subjected to immunofluorescence staining (Anti-IBA1). Quantification analysis of EGFP- and IBA1-double positive cells demonstrated AAV11 has robust transduction capability and approximately 80% specificity in microglia within the spinal cord (Figure 6B and 6C).

**Figure 6.**
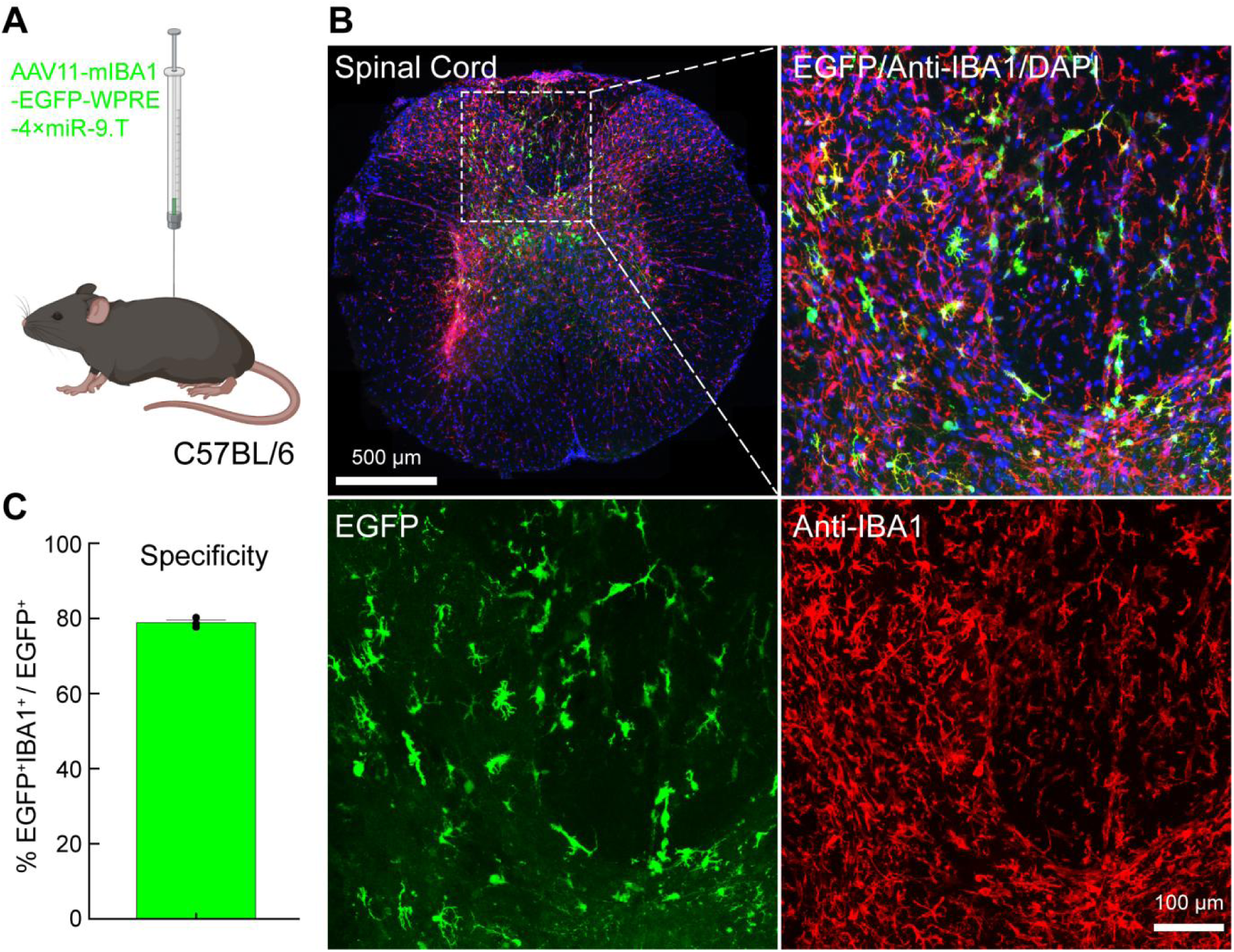
AAV11 efficiently and specifically transduces microglia in the spinal cord. A: Schematic diagram of virus injection. AAV11-mIBA1-EGFP viruses (5 × 10^9^ VG) were injected into spinal cord area of C57BL/6 mice. B: Representative confocal images showing marker expression for AAV11. Scale bar = 500 μm (top left panel), 100 μm (top right panel and bottom 2 panels). C: Quantitative analysis of specificity to microglia for AAV11 (mean ± SEM; *n* = 3 mice per group).

### AAV11 is suitable for sparsely labeling microglia

The elaborate shape of microglia is fundamental to their function in the brain. Microglia have complex processes of change and can undergo morphological changes in different physiological or disease states^44^. Labeling the fine morphology of microglial cells is a prerequisite for understanding their changes in different states and their interaction with other nerve cells^44, 45^. Nevertheless, simple and generalizable labeling methods for studying the fine morphology of microglia within the mammalian brain are still lacking. Here, we injected AAV11 in the hippocampal DG region by reducing the virus injection volume (50 nL), with a dose of 5 × 10^8^ VG (Figure 7A). After three weeks, brain samples were perfused and subjected to immunofluorescence staining (Anti-IBA1). We found that by reducing the injection dose, AAV11 could sparsely label microglia in the hippocampal DG brain area (Figure 7B). By amplifying the signal obtained from 63X laser confocal scanning, the morphology of microglia could be clearly observed (Figure 7C). Therefore, AAV11 can be used for sparse labeling of microglia to analyze their fine structure.

**Figure 7.**
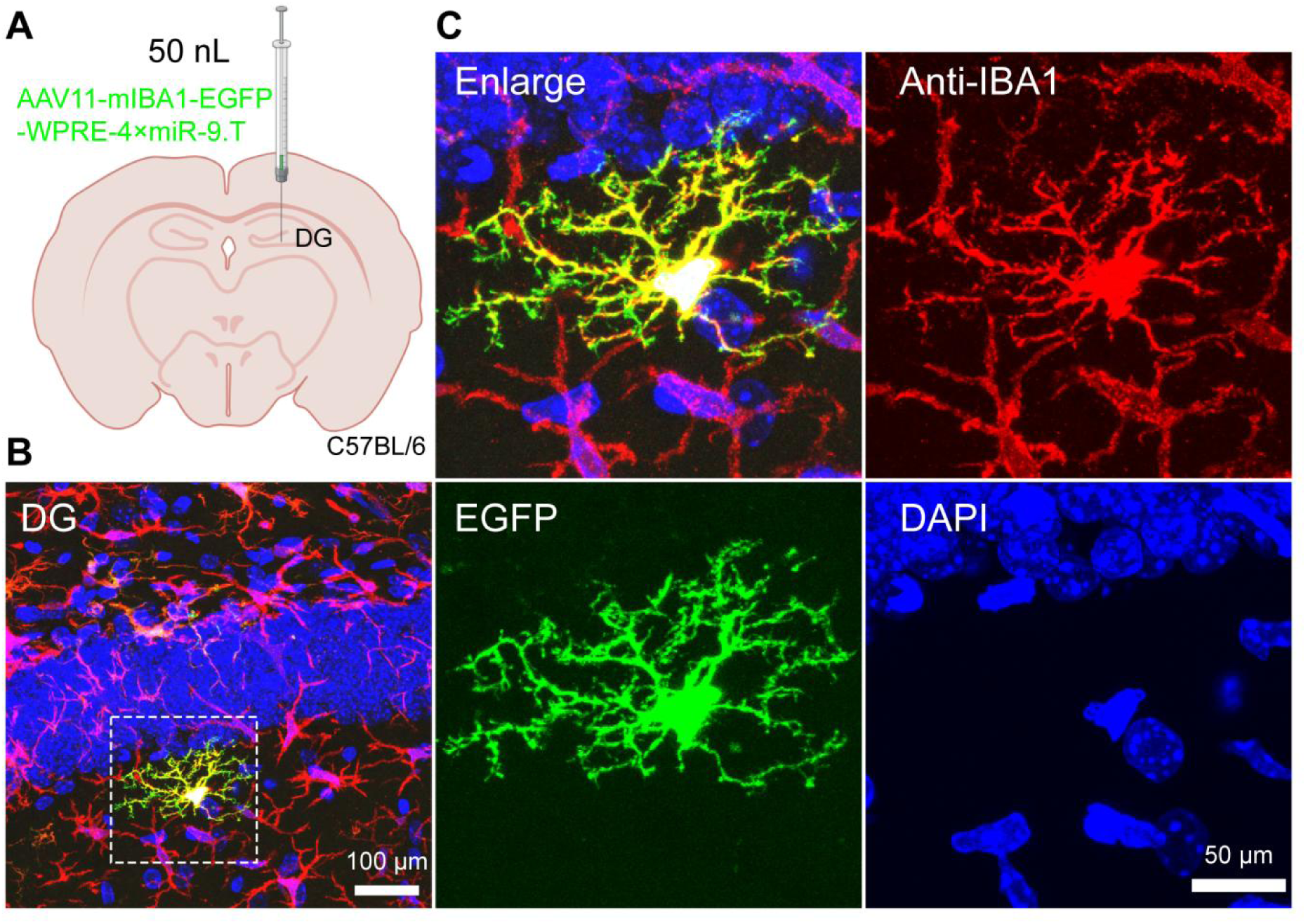
AAV11 sparsely and brightly labels the morphology of microglia. A: Schematic diagram of virus injection. AAV11-mIBA1-EGFP viruses (5 × 10^8^ VG, 50 nL) were injected into DG area of C57BL/6 mice. B: AAV11 sparsely and brightly labeled the morphology of microglia. Scale bar = 100 μm. C: The magnified morphology of microglia labeled by AAV11. Scale bar = 50 μm (right 4 panels).

## Discussion

We achieved microglia-specific transduction in specific brain regions of adult mice through direct injection of AAV11 containing a mouse IBA1 promoter and a set of four tandem sequences for miR-9 targeting. Firstly, we selected natural AAV serotypes including AAV9, AAVhu.32, AAV11 and AAVrh32.33 as the subject of study. AAV11 has previously been reported to efficiently retrograde transduce projection neurons and efficiently transduce astrocytes^46^, indicating its broad tropism and therefore has potential to transduce microglia. AAV9, in prior studies, has been shown to transduce microglia by carrying a mIBA1-specific promoter^38^. AAVhu.32^47^ is highly homologous to AAV9 and may share some of its properties. Additionally, AAVrh32.33 is highly homologous to AAV11 and may also share some of its characteristics. Among these natural serotypes, AAV9 serves as a control for systematically comparing the performance of these AAVs in transducing microglia. We found that AAV11 mediates more specific and efficient transduction of microglia within the CPu. Furthermore, we demonstrated that AAV11 also exhibits high transduction specificity for microglia across distinct brain areas and within the spinal cord. Finally, by reducing the injection dosage, AAV11 could be used for sparse labeling of microglia. This work provides a powerful viral tool for investigating the structure and function of microglia.

The microglial tropism of AAV based tools are mainly determined by the surface proteins in the viral capsid, the cell type specific promoters, and the target sequences of miRNAs that inhibit transgene expression in neurons. Our *in vivo* screens demonstrated that AAV11 armed with the microglia-specific mIBA1 promoter and miRNA-9 targeting sequences exhibited the highest specificity and efficiency in transducing microglia, when compared to other tested AAVs such as AAV9 and AAVhu.32. Previous studies reported that AAVhu.32 tends to transduce neurons with high efficiency^47^, so in accordance with expectation, it performed the worst in microglial cell transduction. This result highlights the importance of the AAV capsid for specific transgene expression in microglia. Moreover, AAV11 demonstrates varying specificity in microglial transduction across different injection regions,, with approximately 97% specificity in the DG, around 90% specificity in the SNr, approximately 70% specificity in the SSp, and approximately 80% specificity in the spinal cord. These different levels of specificity reflect distinct infection patterns of AAV11 in different brain regions. In the DG and SNr, AAV11 may have weaker binding affinity with other cells like neurons. Conversely, in the SSp, AAV11 demonstrates lower microglial transduction specificity, possibly due to the presence of more effective AAV11 binding receptors on these non-microglial cells. More work is needed to demonstrate its efficient transduction mechanism. Of course, these phenomena may also be caused by the performance of promoters and miRNA target sequences in different cells. Therefore, further optimization of microglia-specific promoters and selection of appropriate miRNA target sequences are undoubtedly necessary.

Based on the transduction tropism of microglia, AAV11 can become a new engineering target for developing novel AAVs with more efficient microglial transduction. The advancement of the AAV-MGs benefits from AAV capsid engineering. It is possible to obtain mutant versions of AAV11 through directed evolution^48^ and rational design^23^ to improve microglial targeting ability. For example, we attempted to present the reported functional peptides (such as MG1.1 peptide and MG1.2 peptide)^19^ on the surface of the AAV11 capsid protein to further improve the transduction properties of AAV11 on microglia. In addition, exploring the receptors bound by AAV11 to infect microglia will also provide a reference for the upgrade of microglia-targeting vectors. In summary, a number of AAV11-based tools are expected to be generated soon, which will advance microglia-based biological research and therapeutics.

## Materials and Methods

### Plasmids construction and AAV vector manufacturing

To construct the transfer plasmid pAAV-mIBA1-EGFP-WPRE-4 ⊆miR9.0T-pA, mIBA1 sequence was synthesized and inserted into the pAAV-CD68-EGFP-WPRE-4⊆miR9.0T-pA digested with MluI and SalI restriction endonucleases (Thermo Fisher Scientific, Waltham, MA, USA). AAV packaging plasmids were obtained following the construction method of pAAV2/11^46^. To obtain the pAAV2/hu.32 plasmid, the Cap gene of AAVhu.32 (GeneBank: AY530597.1) was synthesized (Sangon Biotech Co., Ltd., Shanghai, China) and ligated into the pAAV-RC2/1 vector (Addgene, Watertown, MA, USA, 112862). In the principle of point mutation, the Fast Mutagenesis System kit (TransGen Biotech, FM111, China) PCR reaction was applied to construct and amplify pAAV2/rh32.33 plasmid based on pAAV2/11. All primer sequences are placed in Supplementary Table S1. AAV viruses were packaged with the helper plasmid pAd-DeltaF6 (Addgene, Watertown, MA, USA, 112867) and one of the following packaging plasmids: pAAV2/9 (Addgene, Watertown, MA, USA, 112865), pAAV2/hu32, pAAV2/rh32.33 and pAAV2/11^46^.

HEK-293T cells (American Type Culture Collection, Manassas, VA, USA) were cultured in suspension using Balance CD medium (Chuangling Cell-wise, Shanghai, China, CW01001) supplemented with 1% penicillin/streptomycin (BasalMedia, Shanghai, China, S110JV), and kept at 37 ℃ in a 5% CO2 atmosphere. Sixteen hours before the transfection process, these cells were shifted to DMEM (BasalMedia, L110KJ) containing 2% fetal bovine serum (Thermo Fisher Scientific GIBCO, Waltham, MA, USA, 10099141C) and 1% penicillin/streptomycin (BasalMedia, S110JV). Briefly, AAV vectors were generated through a transient transfection method utilizing three plasmids and linear polyethylenimine (Polysciences, Warrington, PA, USA, 24765-1) for the process. At 72 hours following transfection, the viral particles were collected, followed by purification using an iodixanol gradient ultracentrifugation technique, and then concentrated and exchanged into PBS containing 0.001% Pluronic F68 (Thermo Fisher Scientific, Waltham, MA, USA, 24040032) via Amicon® Ultra centrifugal filters (Merck Millipore, Billerica, MA, USA, UFC910024). The titers of the purified recombinant AAVs were determined by quantitative PCR using the iQ SYBR Green Supermix kit (Bio-Rad, Hercules, CA, USA, 1708884) with primer WPRE-F (5’-ATGCCTTTGTATCATGCTATTGCT-3’) and WPRE-R (5’-

CACGGAATTGTCAGTGCCCAA-3’). These viral vectors were then divided into aliquots and stored at -80 ℃ for future use.

### Research animals

All procedures received approval (approval No. APM20026A) from the Animal Care and Use Committee of the Innovation Academy for Precision Measurement Science and Technology, Chinese Academy of Sciences. Adult male C57BL/6 mice (*n* = 27, provided by Hunan SJA Laboratory Animal Company, Changsha, Hunan, China), aged 8–10 weeks, were utilized for the experiments. The mice were housed in a controlled environment under a 12/12-h light/dark cycle within specific pathogen-free facilities. Temperatures were precisely maintained between 22 °C and 24 °C, and humidity levels were kept between 40% and 60%. Both water and food were provided to the animals *ad libitum*.

### Stereotaxic AAV injection

Mice were deeply anaesthetized using 1% pentobarbital intraperitoneally (i.p., 50 mg/kg body weight) and placed in a stereotaxic apparatus (Item: 68025 - stereotaxic apparatus and 68030 - mice adaptor, RWD, China). The injection coordinates were selected according to Paxinos and Franklin’s *The Mouse Brain in Stereotaxic Coordinates*, 4th edition^49^. A small volume of virus was injected into the CPu (relative to bregma: anterior-posterior-axis (AP) +0.80 mm, medial-lateral-axis (ML) ±2.00 mm, and dorsal-ventral-axis (DV) –3.30 mm), DG (relative to bregma: AP –2.15 mm, ML ±1.30 mm, and DV –2.00 mm), SSp (relative to bregma: AP +0.50 mm, ML ±3.00 mm, and DV –2.00 mm) and SNr (relative to bregma: AP –3.1 mm, ML ±1.20 mm, and DV –4.30 mm). Viruses were injected at a rate of 0.03 μL/min using a stereotaxic injector equipped with a pulled glass capillary (Stoelting, Wood Dale, IL, USA, 53311). After the injection was complete, the micropipette was held for an additional 10 minutes before being withdrawn. Animals were allowed to recover from anaesthesia on a heating pad. Three weeks after injection, the mice were sacrificed and brain tissues were collected via transcardic infusion of PBS and 4% paraformaldehyde solution.

### Immunofluorescence and imaging

Brain slices preparation and imaging were performed according to the previously reported methods^50^. The brains were immersed in a 4% paraformaldehyde solution overnight, then dehydrated in a 30% sucrose solution. Subsequently, they were sectioned into 40 μm-thick slices using a microtome (Thermo Fisher Scientific, Waltham, MA, USA), collected in antifreeze solution, and stored at -20 °C until further use. For IBA1 staining, sections were incubated with a primary antibody rabbit anti-IBA1 (1:600, Abcam, Cambridge, MA, USA, ab178846), followed by a secondary antibody donkey anti-rabbit IgG conjugated with Cy3 (1:400, The Jackson Laboratory, Bar Harbor, ME, USA, 711-165-152). After PBS washing, all brain slices mounted on microscope slides were counterstained with DAPI (1:4000, Beyotime, Shanghai, China) and sealed with 70% glycerol. Imaging was conducted using either the Olympus VS120 Slide Scanner microscope (Olympus, Tokyo, Japan) or Leica TCS SP8 confocal microscope (Leica, Wetzlar, Germany).

### Data analysis

Data were analyzed using GraphPad Prism 9.0 (GraphPad Software, La Jolla, CA, USA), average fluorescence intensity was quantified using ImageJ v1.8.0 (National Institutes of Health, Bethesda, MD, USA). Data were presented as means ± standard error of the mean (SEM). All statistical analyses were performed through unpaired two-tailed Student’s *t* tests or one-way ANOVA followed by Tukey’s multiple comparisons test, with significant differences being expressed by the *p* value. Statistical significance was set as *, *p* < 0.05; **, *p* < 0.01; ***, *p* < 0.001 and ****, *p* < 0.0001.

## Data Availability Statement

All datasets for this study are included in the manuscript and the supplementary files.

## Acknowledgments

This work was supported by the National Science and Technology Innovation 2030 Grant (2021ZD0201003), the National Natural Science Foundation of China (31830035, 31771156, 21921004), the Strategic Priority Research Program of the Chinese Academy of Sciences (XDB32030200), the Shenzhen Key Laboratory of Viral Vectors for Biomedicine (ZDSYS20200811142401005), and the Key Laboratory of Quality Control Technology for Virus-Based Therapeutics, Guangdong Provincial Medical Products Administration (2022ZDZ13). Schematic diagrams in the pictures of this article were created with BioRender.com.

## Author Contributions

K.L., and F.X. contributed to the study idea and design; F.X. and J.W. contributed to funding acquisition and resources; N.L., K.L., Y.C., X.S., Z.Z., J.X., G.L. and W.Y. performed the experiments and data acquisition; N.L., and K.L. accomplished data analysis; N.L., K.L., and F.X. drafted the manuscript, and contributed to review and editing. All authors read and approved the final manuscript.

